# Lumen pressure modulation in chicken embryos

**DOI:** 10.1101/2023.12.22.569177

**Authors:** Susannah B.P. McLaren, Fengzhu Xiong

## Abstract

Pressure exerted by fluid contained within a lumen plays a crucial role in the growth, morphogenesis and patterning of epithelial organs. Accurate modulation of lumen pressure in the developing embryo requires sensitive and robust methods that can detect and vary pressure in the range of tens to hundreds of Pascals (Pa). Here we describe a simple, cost-effective method for setting up a pressure modulation apparatus combining a high-sensitivity pressure sensor and a water column whose height can be finely tuned. We demonstrate lumen pressure control using the developing brain of early chicken embryos.

## 1 Introduction

A fluid-filled lumen bounded by an epithelium is a common feature in development and forms the basis of many early organs. Hydrostatic pressure within such a lumen can drive organ expansion [1] and impacts multiple biological processes such as changes in tissue mechanics [2], cell movement [3] and cell rearrangement [4]. Pressure increase in the lumen of the developing zebrafish inner ear generates stress within the surrounding epithelium and subsequent organ expansion [1]. In the developing mouse embryo, an increase in lumen pressure leads to mechanical stretching of epithelial cells that line the lumen [2] and guides the positioning of the first axis of symmetry in the embryo [5]. The brain of chicken embryos folds abnormally when the lumen is punctured, suggesting that a positive internal pressure is required to maintain neural tissue shape and brain volume [6].

The ability to robustly modulate pressure within a lumen will aid our understanding of the role hydrostatic pressure plays in biological processes. Here we describe a method utilising easily obtainable and simple components, drawing inspiration from previous studies [1, 2, 5, 7]. Hydrostatic pressure control is achieved by combining a high-precision sensor and a water column. This closed fluid system links to a needle that punctures the target lumen, equating its pressure to that of the adjustable water column. Using the developing chicken embryo brain as an example, we show that the apparatus causes expansion and collapse of the epithelial lumen. This method can be easily adapted for other model systems and can also be used for different purposes such as pipette aspiration.

## 2 Materials

### 2.1 Pressure probe and modulation apparatus

A pressure sensor, such as the Honeywell HSCDANT001PGSA3, is required to enable a live readout of the hydrostatic pressure. For sensor output, a microcontroller, such as an Arduino Uno, connected to a computer via USB can be used. An electronics breadboard and jumper wires are used for circuit connections. Needles for piercing biological cavities are made by pulling glass capillary tubes with a micropipette puller, for instance, one from Sutter Instrument. Various tubing is necessary for fluid handling. We used such Tygon tubing with 1-6mm inner diameters. The system incorporates a T-junction with a fluid splitter valve to manage fluid flow. A 5 ml syringe is used for filling the tubing, needle, and sensor port with phosphate-buffered saline (PBS). Additionally, a larger syringe (60ml) is used for creating the water column. For sample observation, a micromanipulator, such as the M3301-M3 from World Precision Instruments, controls needle insertion into the sample, and a linear stage, for example, the LNR50M/M from Thorlabs, is used for fine adjustment of the height of the water column. Finally, a stereomicroscope with a camera, like the Nikon SMZ745, is utilized for monitoring the sample.

### 2.2 *Ex ovo* chicken embryo culture

Early-stage chicken embryos were obtained from fertilized chicken (*Gallus gallus domesticus*). Eggs are incubated in a ventilated and humidified environment (∼40% humidity) maintained at a temperature range of 37.5-38.5°C. For microsurgical procedures, tools such as forceps, surgical scissors, microcapillary needles, and a scalpel blade are required. Cardboard egg holders, typically received with the egg shipment, are used for stabilizing the eggs. Two sizes of Petri dishes, 35mm and 100mm, are needed for various stages of the process. Additionally, 2cm x 2cm pieces of filter paper, e.g. Whatman brand, with two partially overlapping holes (each 0.5cm in size) punctured using a paper puncher, are prepared. A small metal keyring is used to prevent the embryo from floating in phosphate-buffered saline (PBS). An egg waste bag is kept handy for disposal purposes. For ex ovo culture plates, reagents and tools such as Bacto-Agar, a 20% glucose solution (filtered), and 5M NaCl (autoclaved) are essential. Other necessary equipment includes a water bath set to 55°C, a microwave, and a magnetic stirrer.

## 3 Methods

### 3.1 Apparatus assembly

To assemble the pressure modulation apparatus, we first tailored 3 pieces of tubing of appropriate inner diameters and lengths according to the layout of equipment (e.g. Figure 1) and their respective connections. We used a pair of sharp forceps to trim the insertion needle tip to the required diameter (See *Note 1*) and filled the insertion needle with PBS using a 5ml syringe with an attached needle fitting. Each piece of tubing and the T junction was filled with PBS and the first piece of tubing was connected to the insertion needle and then to the 3-way T junction. We then gently injected a small amount of PBS into the port of the pressure sensor to replace the air in the port whilst ensuring that the electrical pins of the sensor stayed dry. After connecting the second PBS-filled piece of tubing to the pressure sensor port, we then connected it to the opposite side of the 3-way T junction, taking care to avoid trapping of air bubbles (See *Notes 2 and 3*). The plunger from a 60ml syringe was removed and the column was filled with a desired amount of PBS, whilst covering the open end of the syringe. The third piece of PBS-filled tubing was then connected to the PBS column and to the 3-way T junction (whilst keeping the connection closed to the PBS column), at right angles to the other tubes. To enable precise adjustment of the PBS column’s height, we connected the PBS column to an adjustable linear stage, with which the height of the column can be finely controlled. Finally, the insertion needle was mounted to the movable arm of the micromanipulator which was used to position the needle tip so that it was visible and in-focus when viewed using the stereomicroscope and camera. The needle tip should lie at a similar height as the sensor when inside the sample.

**Figure 1.**
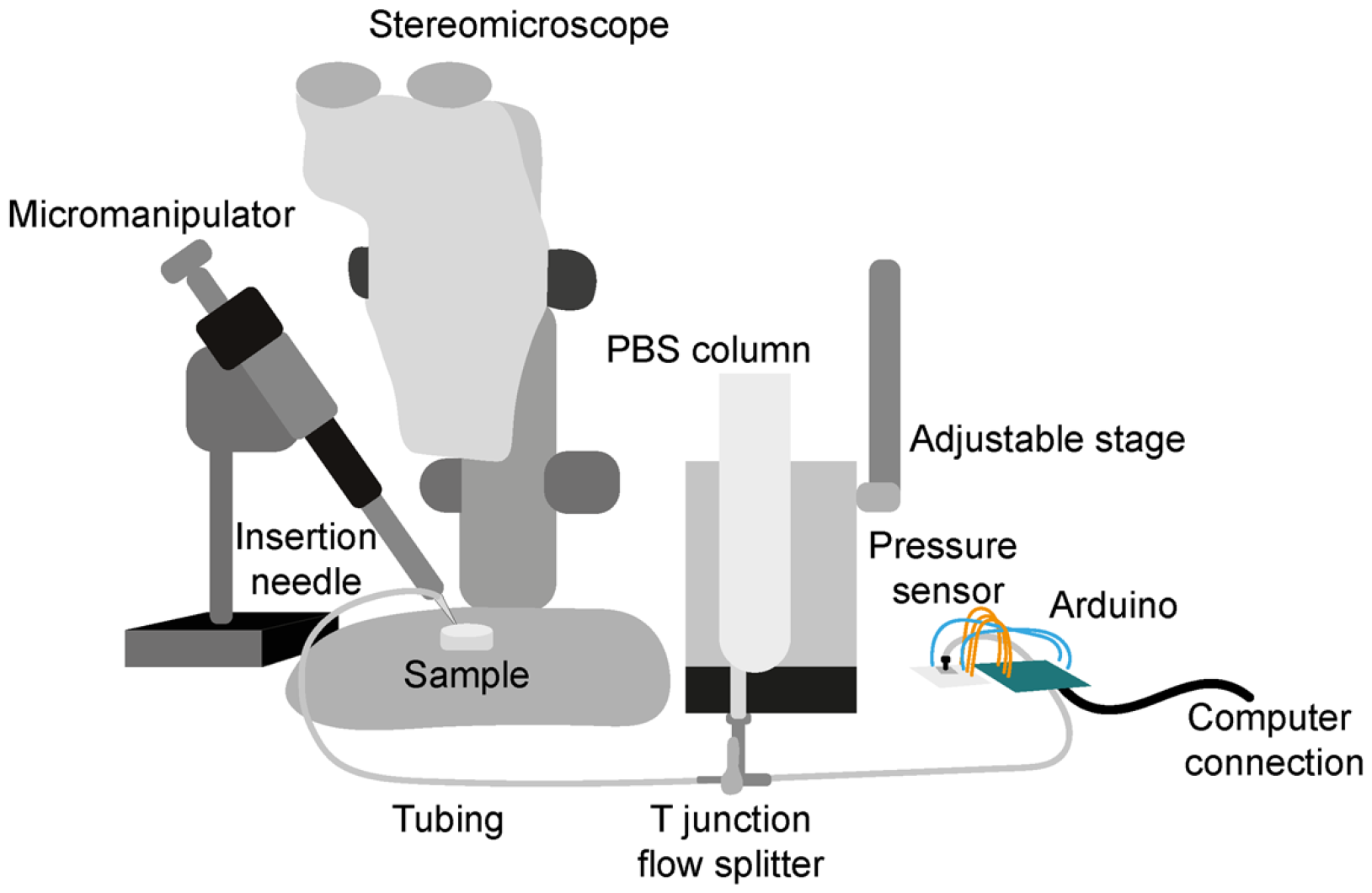
Illustration depicting pressure modulation set-up. The set-up can be assembled on a standard lab bench and the arrangement of components customised to the experimenters needs.

### 3.2 Electronic connections and software set-up

Pressure recordings were visualised and saved in real time by connecting the pressure sensor to a computer via a microcontroller board. First, we connected the pressure sensor to a breadboard and used jumper wires to connect each sensor pin to the Arduino Uno as required for the sensor output type. As an example, the pin connections are given for a SPI output sensor (Honeywell HSCDANT001PGSA3): Sensor Pin 1 – GND; Sensor Pin 2 – Voltage supply (use 3.3V); Sensor Pin 3 – MISO (Arduino Pin 12); Sensor Pin 4 – SCLK (Arduino Pin 13); Sensor Pin 5 – SS (Arduino Pin 10). The Arduino Uno was then connected to a computer using a USB cable. We used the Arduino IDE user interface and downloaded the HoneywellTruStability SPI library (Erik Werner - https://github.com/huilab/HoneywellTruStabilitySPI) to read from the Honeywell pressure sensor. The “HSCPressureSensorTest” code was opened in Arduino IDE and modified accordingly to fit the specific sensor. In our case a 0-1PSI (0-6.9kPa) gage pressure sensor was used. Once modified, we uploaded the Arduino code to the pressure sensor. For initial tests, we used the ‘serial read’ or the ‘serial monitor’ in the Arduino IDE to print or plot the pressure readings, respectively. See also *Note 4*. Code enabling reading, writing to csv and plotting of the data with Python can be found at: https://github.com/susie-m/lumen_pressure_reading

### 3.3 Calibration

We submerged the insertion needle in the water/PBS column and used the hydrostatic pressure at different water depths to calibrate the probe. To do this, we fixed the insertion needle in place so that it was suspended in such a way that its position remained fixed when the column was moved vertically by the adjustable stage. We adjusted the height of the column, and so the depth of the needle tip in the PBS, by moving the stage up or down in fixed intervals e.g. 1mm and recorded the pressure at each depth, checking that the pressure change corresponded to that expected for the depth change. A pressure difference of approximately 9.8Pa is expected for a change in depth of 1mm H_2_O (Figure 2). A linear relationship was observed between steps in depth and the pressure reading.

**Figure 2.**
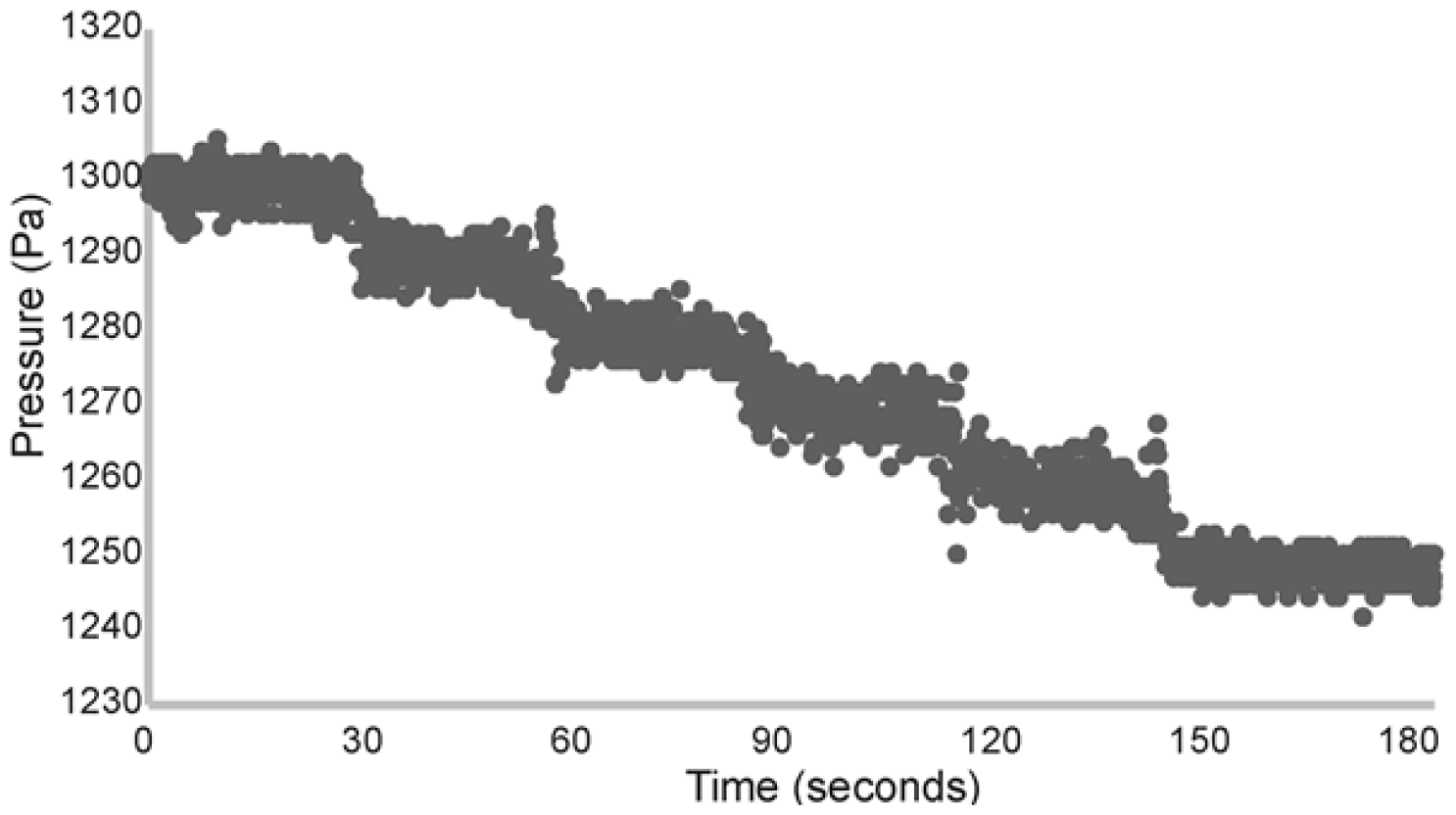
Example calibration recording. Pressure profile corresponding to the depth of the insertion needle during calibration. The depth of the needle tip in PBS was decreased by 1mm approximately every 30 seconds by manually adjusting the height of the PBS column using an adjustable stage. This corresponded to a pressure decrease of approximately 10Pa every 30 seconds. Pressure readings were sampled every 0.1 seconds.

### 3.4 Embryo preparation

#### 35mm *ex ovo* culture dishes were prepared as follows

∼150ml albumen (egg white) was collected and a magnetic stirrer used to beat the albumen for ∼15 minutes at medium speed. Foam was removed and 100ml of clear albumen was collected in a beaker. 1.5ml of 20% Glucose was added to the beaker and the beaker was placed in a 55°C water bath. 0.6g Bacto Agar gel was dissolved in 100ml water in a glass beaker in the microwave and 2.5ml 5M NaCl was added following this. The gel mixture was allowed to cool until the glass could be comfortably touched. Whilst still warm, the gel mixture was poured into the albumen beaker in the 55°C water bath and the solutions were mixed. 2ml of the combined mixture was pipetted into each 35mm petri dish and the gel was left to set. Dishes were kept in 4°C for storage.

#### Embryos were prepared as follows

Fresh (within 10 days of receipt, stored in 14°C fridge) fertilized chicken eggs were incubated at 37°C for 64-72 hours. An embryology workspace was prepared so that microsurgical tools, one empty 100ml petri dish, one PBS filled 100ml petri dish, a bag for egg waste, and pre-punctured filter paper pieces were available. Culture dishes were placed in an incubator to warm up 10 minutes prior to starting embryo culture. A cardboard egg box was used to hold each egg in place whilst drilling a small hole into the side of the egg using the sharp point of a scalpel blade until a crack appeared. The egg was gently prised open, allowing the yolk with the embryo on top to drop into the petri dish, with the embryo facing upwards. A piece of filter paper soaked in PBS was used to gently remove albumen from around the embryo by sweeping it to the side. A hole-punched small piece of filter paper was placed over the embryo, so that the embryo AP axis lay in the centre of the long axis of the holes. After gently pressing down on the filter paper with forceps sharp dissection scissors were used to cut through the yolk following the outer edge of the filter paper. The filter paper with embryo attached was peeled away from the yolk using forceps, washed briefly in the PBS petri dish and placed on a pre-warmed culture dish. Cultures were kept in a humidified box in the incubator.

When ready to begin pressure modulation, the culture dish was placed in the field of view on the stereomicroscope and a sharp microcapillary needle was used to gently remove the vitelline membrane from the region targeted for pressure modulation. A small metal ring was placed on top of the filter paper surrounding the embryo so that the embryo lies in the middle of the ring and pre-warmed PBS was added to submerge the embryo.

## 4 Results

### 4.1 Pressure modulation

Pressure modulation was achieved by adjusting the height of the PBS column so that a difference between the level of the PBS and the insertion needle tip was created (Figure 3a and b). Pressure was increased by raising the height of the PBS level above the level of the insertion needle tip or decreased by lowering the column.

**Figure 3.**
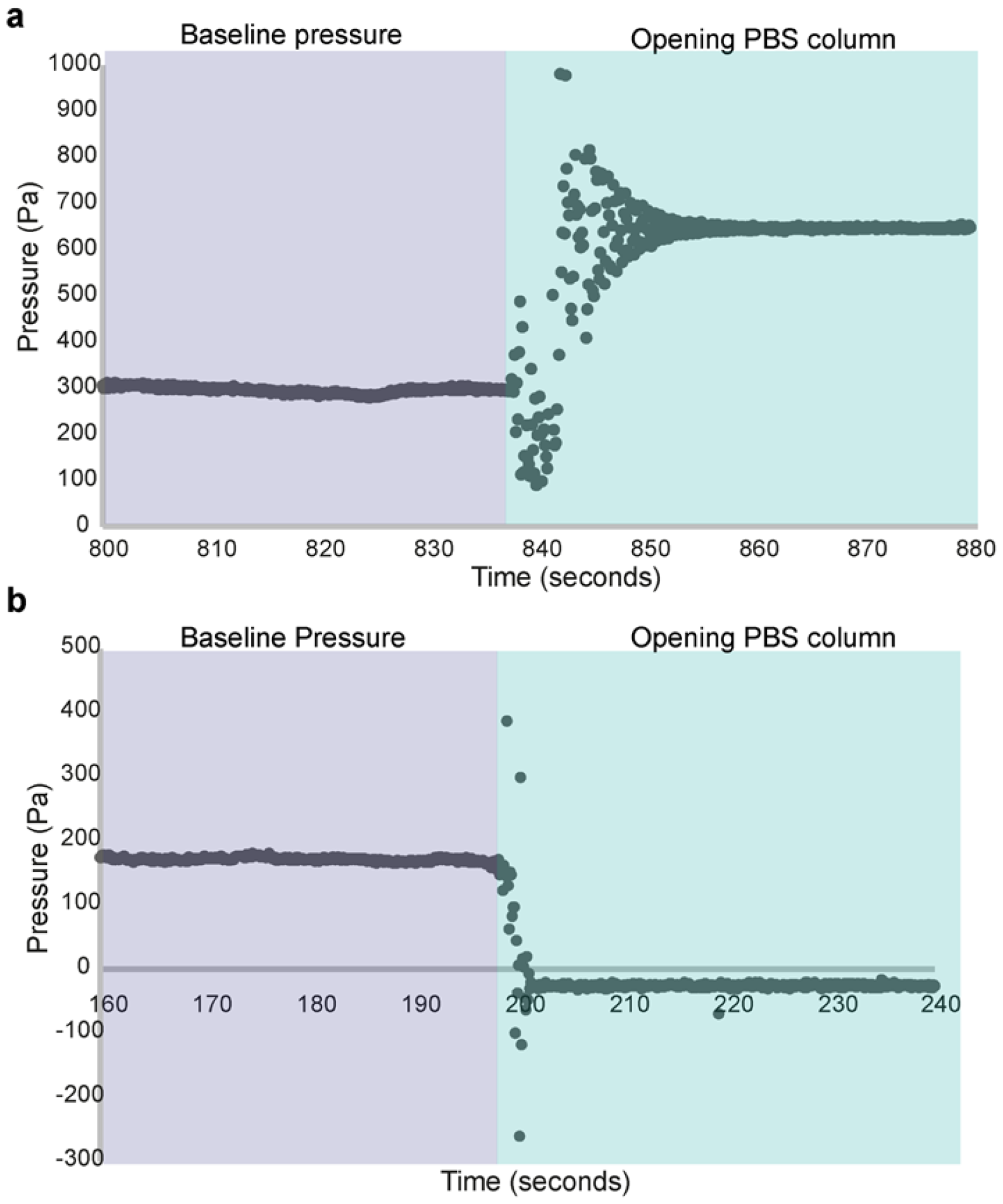
Example profiles for pressure increase and decrease. (a). Pressure profile observed whilst insertion needle is inside brain (baseline) and after the channel to PBS column has been opened when PBS level is raised above the needle tip. In this example pressure was increased by approximately 340Pa. Pressure readings were sampled every 0.1 seconds. (b). Pressure profile observed whilst insertion needle is inside brain (baseline) and after the channel to PBS column has been opened when PBS level is lowered below the needle tip. In this example pressure was decreased by approximately 140Pa. Pressure readings were sampled every 0.1 seconds.

Lumen pressure was increased by initially setting the PBS column to the desired height above the insertion needle whilst the connection to the PBS column was closed. The needle was then inserted into the PBS in the dish next to the embryo. The pressure readings should be stable when the needle is held stationary (See *Note 5*). To test the efficacy of the pressure modulation set-up we sought to increase and decrease pressure in the neural tube lumen. We targeted the anterior embryonic brain where the lumen is relatively wide and inserted the needle into the lumen in this region. Once the lumen had been successfully punctured, the needle was maintained in its current position and the pressure profile recording was observed. The connection to the PBS column was then opened to increase lumen pressure (Figure 3a). Lumen inflation was observed on successful puncture and pressure increase (Figure 4). To decrease lumen pressure the PBS column was lowered below the level of the needle tip, rather than raised. This led to observable deflation of the lumen and collapse of the brain structure (Figure 3b and Figure 4).

**Figure 4.**
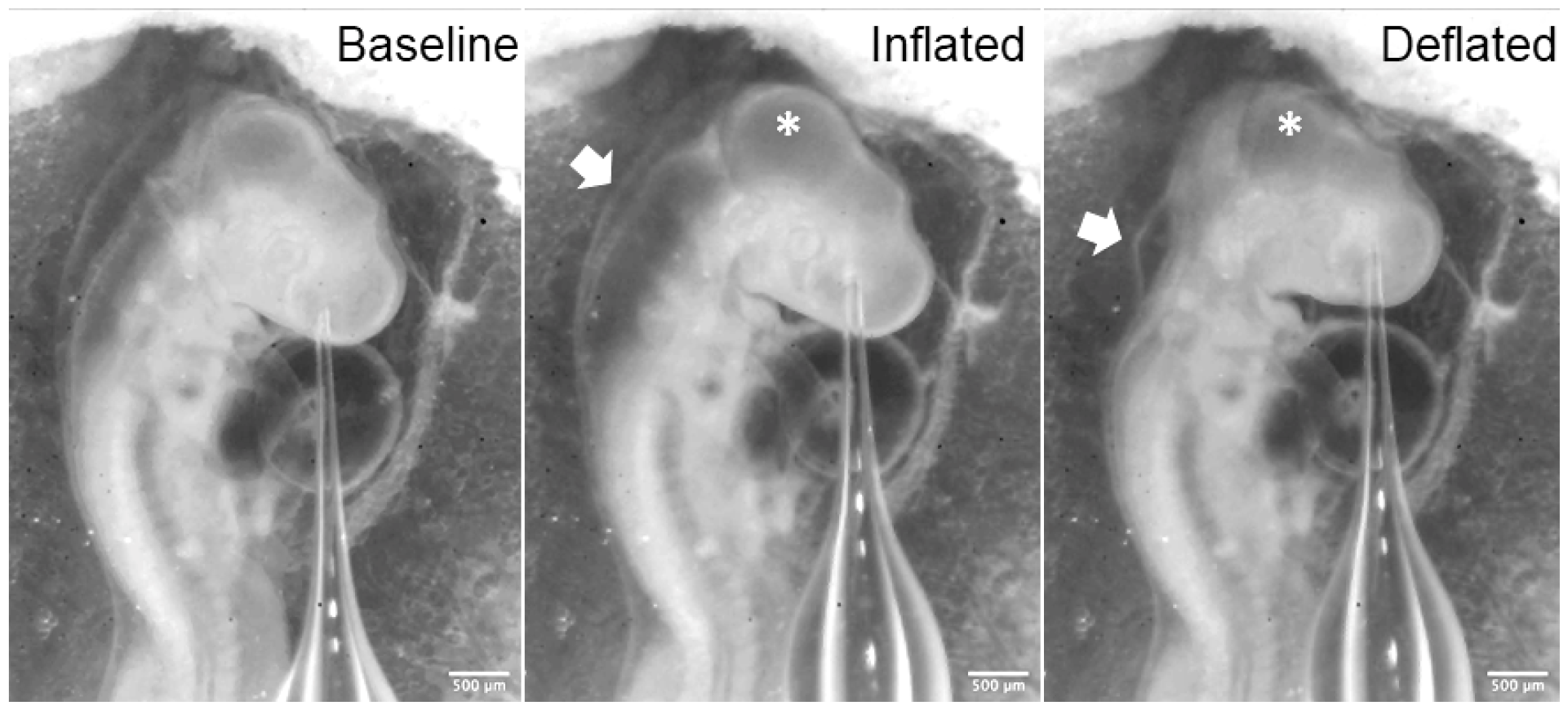
Lumen inflation and deflation. Example of brain inflation in a HH16 stage chicken embryo with needle inserted into forebrain. White arrow points to the hindbrain area. Asterisk marks the midbrain. Clear inflation and deflation can be observed in these structures. Scale bars are 500μm.

Together, these results demonstrate the ability of this pressure modulation apparatus to increase and decrease pressure in a controlled way in the neural tube lumen of chicken embryos. This setup offers a versatile and customisable way to modulate hydrostatic pressure in a range of biological systems and enables investigation of the role of fluid pressure in the development of tissues and organs with a fluid-filled lumen.

## 5 Notes

1. The diameter of the needle tip often needs to be adjusted according to the type and status of the sample to prevent clogging (increasing diameter) whilst still allowing for puncturing and a good seal between the needle tip and lumen tissue. It is useful to open the needle tip at an angle to create a slanted cross-section to help tissue puncturing.
2. It is important that the seals between each piece of tubing and the connected apparatus are as tight as possible, preventing and leakage of PBS and the entry of air into the system.
3. Before starting modulation, check that air bubbles are not present in the connected set up. If present, air bubbles can be flushed out by opening the connection to the PBS column and allowing PBS to drain through the system when the tubes are disconnected from the sensor and needle. Alternatively disconnect each piece of tubing, remove air bubbles using the syringe, re-fill with PBS, and reconnect.
4. Customization and further upgrades (optional). Serial data from the Arduino can be monitored and recorded using custom scripts. It is also possible to synchronize the frames of the camera stream with pressure reads by programming the Arduino to trigger the camera. To implement precise pressure control for long-term experiments, a motorized stage can be used to replace the adjustable linear stage to hold the water/PBS column. Then program the sensor readings to feedback to the control of the motorized stage. Incubation environment can be introduced by moving the needle part of the system to an existing microscope with environmental chamber, or by adding a heating box/stage around the sample with a transparent top cover.
5. Sharp bends in the tubing should be avoided and tubing should be positioned so that it stays as stationary as possible as small fluctuations in tubing height can lead to unstable pressure recordings.

## Acknowledgement

This work is supported by a Wellcome Trust / Royal Society award (215439/Z/19/Z).

